# Electron microscopy 3-dimensional segmentation and quantification of axonal dispersion and diameter distribution in mouse brain corpus callosum

**DOI:** 10.1101/357491

**Authors:** Hong-Hsi Lee, Katarina Yaros, Jelle Veraart, Jasmine Pathan, Feng-Xia Liang, Sungheon G. Kim, Dmitry S. Novikov, Els Fieremans

## Abstract

To model the diffusion MRI signal in brain white matter, general assumptions have been made about the microstructural properties of axonal fiber bundles, such as the axonal shape and the fiber orientation dispersion. In particular, axons are modeled by perfectly circular cylinders with no diameter variation within each axon, and their directions obey a specific orientation distribution. However, these assumptions have not been validated by histology in 3-dimensional high-resolution neural tissue. Here, we reconstructed sequential scanning electron microscopy images in mouse brain corpus callosum, and introduced a semi-automatic random-walker (RaW) based algorithm to rapidly segment individual intra-axonal spaces and myelin sheaths of myelinated axons. Confirmed with a conventional machine-learning-based interactive segmentation method, our semi-automatic algorithm is reliable and less time-consuming. Based on the segmentation, we calculated histological estimates of size-related (e.g., inner axonal diameter, g-ratio) and orientation-related (e.g., Fiber orientation distribution and its rotational invariants, dispersion angle) quantities, and simulated how these quantities would be observed in actual diffusion MRI experiments by considering diffusion time-dependence. The reported dispersion angle is consistent with previous 2-dimensional histology studies and diffusion MRI measurements, though the reported diameter is larger than those in other mouse brain studies. Our results show that the orientation-related metrics have negligible diffusion time-dependence; however, inner axonal diameters demonstrate a non-trivial time-dependence at diffusion times typical for clinical and preclinical use. In other words, the fiber dispersion estimated by diffusion MRI modeling is relatively independent, while the "apparent" axonal size estimated by axonal diameter mapping potentially depends on experimental MRI settings.

## Introduction

Diffusion MRI (dMRI) is a noninvasive imaging modality that, by measuring the random motion of water molecules at clinically accessible diffusion times *(t* ~ 50 ms), is sensitive to a length scale of about ten micrometer, comparable to cell sizes. Several tissue models for dMRI have been proposed to specifically probe the neuronal microstructure, in order to estimate the axon orientation dispersion (Jespersen et al. 2017; Novikov et al. 2018b; Reisert et al. 2017; Ronen et al. 2014; Schilling et al. 2018; Tariq et al. 2016; Zhang et al. 2012) and the axonal diameter distribution (Alexander et al. 2010; Assaf et al. 2008; Barazany et al. 2009; Benjamini et al. 2016; Duval et al. 2015; Komlosh et al. 2013). To enable dMRI modeling, assumptions have been made, which so far have not been fully validated by comparing against histology. In particular, axons are assumed to be circular cylinders with no diameter variations along each axon (Alexander et al. 2010; Assaf and Basser 2005; Assaf et al. 2008). This widely adopted assumption contradicts histological observations of neurite beadings (varicosities) (Baron et al. 2015; Budde and Frank 2010; Li and Murphy 2008; Shepherd et al. 2002; Tang-Schomer et al. 2012), and needs to be further evaluated when modeling the orientation dispersion and the axonal diameter distribution, particularly with respect to their dependency on experimental settings such as the diffusion time *t*.

To validate dMRI models in the brain white matter (WM) against histology, previous studies reported multiple tissue parameters, including the fiber orientation distribution (FOD) (Grussu et al. 2016; Mollink et al. 2017; Ronen et al. 2014; Schilling et al. 2016; Schilling et al. 2018), axon dispersion angle (Ronen et al. 2014), inner axonal diameter (Abdollahzadeh et al. 2017; Aboitiz et al. 1992; Caminiti et al. 2009; Kleinnijenhuis et al. 2017; Liewald et al. 2014; Mason et al. 2001; West et al. 2015), and g-ratio (Abdollahzadeh et al. 2017; Kleinnijenhuis et al. 2017; Mason et al. 2001; Stikov et al. 2015; West et al. 2015; West et al. 2016; Yang et al. 2016). In histology, size-related quantities, e.g., diameter and g-ratio, were estimated either by 2d (Aboitiz et al. 1992; Caminiti et al. 2009; Liewald et al. 2014; Mason et al. 2001; West et al. 2015) or 3d high-resolution electron microscopy (EM) images (Abdollahzadeh et al. 2017; Kleinnijenhuis et al. 2017). In contrast, the orientation-related metrics, e.g., FOD and dispersion angle, were evaluated either by 2d low-resolution polarized light images (4 μm/pixel, in-plane) (Mollink et al. 2017) and light microscopy images (1-1.6 μm/pixel, in-plane) (Grussu et al. 2016; Ronen et al. 2014), or by 3d moderate-resolution confocal microscopy images (0.42 μm/slice, through plane) (Schilling et al. 2016; Schilling et al. 2018). Although the tissue anisotropy index has been evaluated on 3d EM images (Salo et al. 2018), retrieving other orientation metrics, e.g., FOD, rotational invariant and dispersion angle, from 3d high-resolution EM images (≤ 0.1 μm/slice, through plane) has not yet been attempted.

Furthermore, definitions of tissue parameters differ between studies, both in histology and MRI, and need to be clarified before use. In particular, the orientation dispersion of axon bundles could be summarized by (1) the standard deviation (SD) of dispersion angles projected on a 2d plane (Ronen et al. 2014), by (2) rotational invariants and the root-mean-square (rms) of the dispersion angle’s cosine for spherical harmonics (SH)-based methods (Jespersen et al. 2017; Novikov et al. 2018b; Reisert et al. 2017), or by (3) the normalized orientation dispersion index (ODI) of specific orientation distributions (e.g., Watson or Bingham distributions) (Schilling et al. 2018; Tariq et al. 2016; Zhang et al. 2012). In this study, we focus on the first two definitions to avoid introducing further assumptions on the axon dispersion.

Similarly, axonal diameters have been estimated using various methods. To avoid overestimation of the inner axonal diameter caused by obliquely sliced axons, most of the 2d histological studies measured the inscribed circle diameter as the diameter estimate (Aboitiz et al. 1992; Caminiti et al. 2009; Liewald et al. 2014). Other histological studies adopted an equivalent circle diameter calculated by the cross-sectional area (West et al. 2015) or the short axis length of a fitted ellipse (Abdollahzadeh et al. 2017). In this study, we focus on the equivalent circle diameter since (1) the 3d axon structure is fully reconstructed and free from problems of oblique cross-sections, and (2) contours of the intra-axonal space and the myelin sheath might be different, leading to unreliable estimates of the g-ratio (ratio of inner to outer diameter) when using other definitions.

Aside from ambiguous definitions of microstructural features, comparison between dMRI and histology depends on the experimental settings of the dMRI experiment. Indeed, the diffusion process can be approximately understood as a coarse-graining process, which is equivalent to smoothing the tissue microstructure (Novikov et al. 2014) using a kernel of a width commensurate with the diffusion length 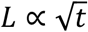. In other words, the diffusion times applied in dMRI measurements potentially affect the metric estimations between studies using different acquisition parameters, and have to be carefully accounted for.

So far, many EM segmentation software tools have been released for segmenting gray matter images that use semi-automatic analysis with an interactive proofreading interface (Benjamini et al. 2016; Dorkenwald et al. 2017; Jeong et al. 2009; Kaynig et al. 2015; Sommer et al. 2011). Such methods require abundant training data, for which generation is very labor-intensive and time-consuming. In WM, however, appropriate EM segmentation methods are still limited (Abdollahzadeh et al. 2017; Kleinnijenhuis et al. 2017; Zaimi et al. 2018). To segment myelinated axons within acceptable processing time, we propose here a semi-automatic segmentation algorithm depending on diffusion trajectories obtained by random-hopping on a cubic lattice bounded by a binary myelin mask, as a simplified version of the seeded-region-growing method (Abdollahzadeh et al. 2017; Adams and Bischof 1994). Our segmentation method was further validated by comparing against the interactive Carving function in ilastik (Sommer et al. 2011).

Here, by analyzing segmented myelinated axons, we calculated both orientation-related and size-related axonal features based on definitions from either histology or MRI experiment, where the effect of varying diffusion time was simulated by applying a corresponding smoothing kernel, and demonstrated a non-trivial discrepancy between estimated microstructural characteristics from both definitions. In particular, we demonstrated by 3*d* high-resolution EM segmentation, for the first time, the influence of the diffusion time-dependence on the orientation dispersion and the inner axon diameter.

## Materials and Methods

All procedures performed in studies involving animals were in accordance with the ethical standards of the institution or practice at which the studies were conducted. This article does not contain any studies with human participants performed by any of the authors.

### Animals and image acquisition

A female 8-week-old C57BL/6 mouse was perfused trans-cardiacally using a fixative solution of 4% PFA, 2.5% glutaraldehyde, and 0.1M sucrose in 0.1M phosphate buffer (PB, pH 7.4). The genu of corpus callosum was later excised from the dissected brain, and the tissue was fixed in the same fixative solution, followed by a PB containing 2% OsO4 and 1.5% potassium ferrocyanide for 1 hour. The tissue was then stained with 1% thiocarbohydrazide (Electron Microscopy Scientices, EMS, PA) for 20min, 2% osmium tetroxide for 30min, and 1% aqueous uranyl acetate at 4°C overnight. An *En Bloc* lead staining was performed at 60°C for 30 min to enhance membrane contrast. The brain sample was dehydrated in alcohol and acetone, and embedded in Durcupan ACM resin (EMS, PA (Wilke et al. 2013)). The tissue sample was analyzed with a scanning electron microscope (SEM) (Zeiss Gemini 300 SEM with 3View), and 401 consecutive images of 6000 x 8000 pixels were acquired, representing a volume of 36 x 48 x 40.1 μm^3^ with a resolution of 6 x 6 x 100 nm^3^.

### Image processing and axon segmentation by Random-Walker algorithm (RaW)

We down-sampled the image to a resolution of 24 x 24 x 100 nm^3^ using Lanczos interpolation to lower the computational cost without compromising the segmentation accuracy. Further, we corrected the geometric distortion in slices disagreeing with the interpolation estimated from adjacent slices, by using a non-linear deformation calculated with optical flow estimation (Sun et al. 2010; Sun et al. 2014), and selected a subset of 200 slices (36 x 48 x 20 μm^3^ in volume) to rule out slices with intractable distortions (Fig. 1a).

**Fig. 1.**
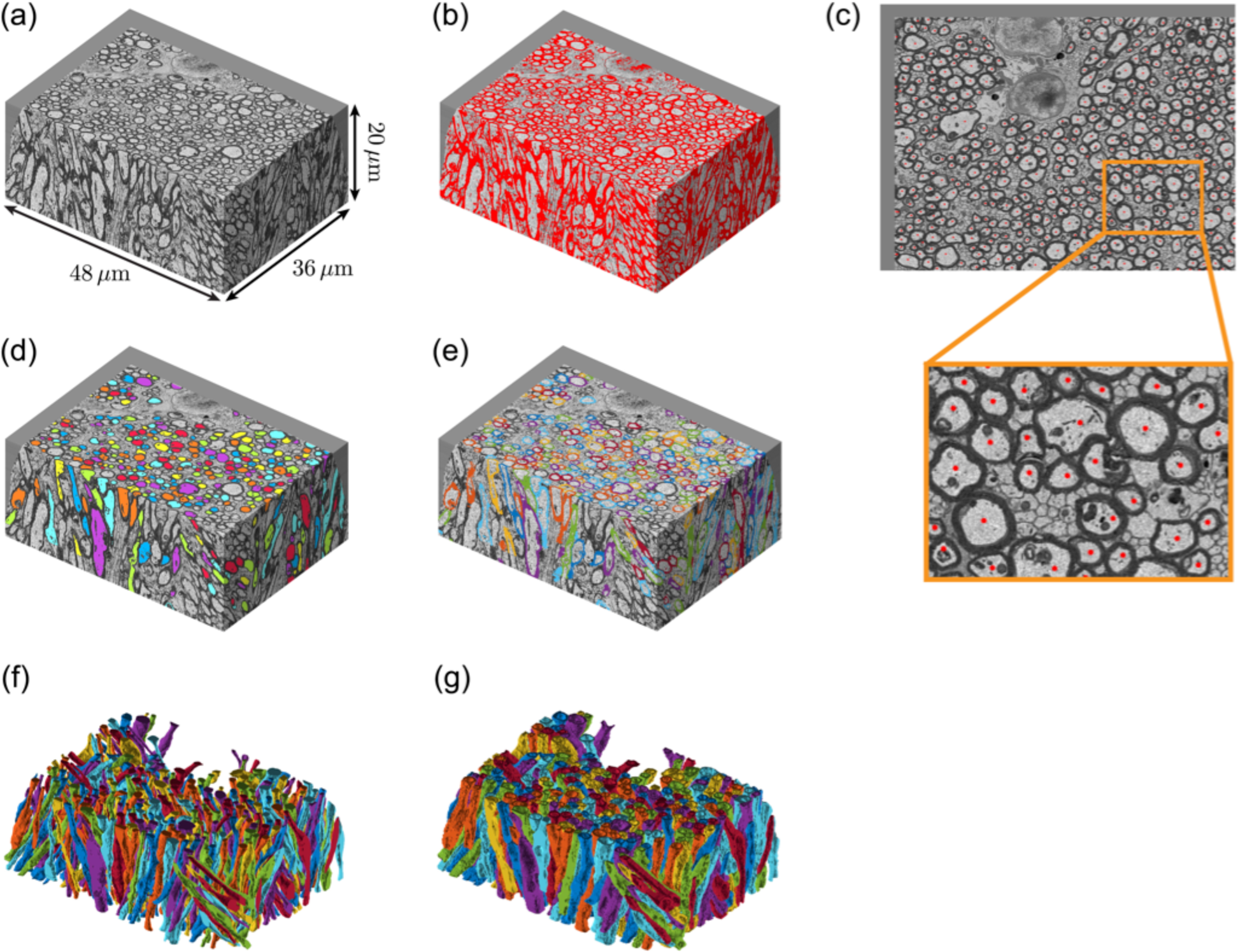
The semi-automatic IAS segmentation pipeline. (a) A tissue sample of genu in CC, in a volume of 36 x 48 x 20 μm^3^, was acquired by sequential SEM. (b) The myelin mask (red) was calculated by intensity thresholding with a bi-Gaussian model for further segmentation of the intra-axonal space (IAS). (c) Seeds (red dots) for random diffusion grid-hopping process were assigned manually over one central slice (451 seeds). The random-hopping trajectory was bounded by the myelin mask in (b). (d) IAS (colors) was filled by all random-hopping trajectories (316 segmented IAS). The IAS from axons with leaky myelin mask has been excluded by proofreading. (e) The individual myelin sheath (colors) is the overlap of the myelin mask and the expanded IAS dilated by ≤ 0.4 μm. Touching myelin sheaths of adjacent axons are separated based on a non-weighted watershed algorithm. (f-g) By transforming each segmented IAS and individual myelin sheath into polyhedrons, it is feasible to perform numerical simulations in such 3d realistic microstructure

To semi-automatically segment the intra-axonal space (IAS) of myelinated axons, we employed a random-hopping diffusion process, dubbed Random-Walker algorithm (RaW), as a simplified seeded-region-growing algorithm (Adams and Bischof 1994) applied on a binary mask: We manually seeded an initial position per axon within the central slice (451 seeds in Fig. 1c), and filled the IAS by diffusion trajectories, obtained by random-hopping on a cubic lattice, of 4000 particles per axon with 640,000 steps. The diffusion trajectory is confined by a myelin mask (Fig. 1b) obtained by fitting and thresholding the EM image intensity histogram with a bi-Gaussian model. Segmented axons with an imperfect myelin mask were deleted by proofreading, resulting in an IAS mask of 316 segmented axons (~ 51,000 axon segments). The IAS segmentation was then completed by automatically seeding within the previously generated diffusion trajectories confined by the non-leaky myelin mask (Fig. 1d). The seeding density is a seed per ten slices for each axon, filled with 10,000 particles per seed with 40,000 steps. The IAS mask was downsampled into (100 nm)^3^-resolution to further analyze axon geometries, e.g., fiber dispersion, axonal diameter, myelin thickness, and g-ratio.

All the processing was implemented in Matlab and accelerated by parallel computation with 12 CPU cores. Total processing time, including the manual seeding, numerical computations and the proofreading, was 1-2 days.

### Conventional axon segmentation (ilastik, Carving)

To compare RaW with a conventional segmentation method, the IAS of 100 selected axons was manually carved by K.Y. and J.P., using the ilastik software package (Sommer et al. 2011). Seeded watershed segmentation was performed by ilastik with following settings:

1. Each 2*d* image (resolution = 24 x 24 nm^2^) was smoothed by a kernel with a size of 1.6 pixel (38.4 nm), and was filtered to enhance edges, e.g., cell membrane, mitochondria and myelin sheath.
2. Markers inside the IAS (object seeds) and outside the IAS (background seeds) were manually assigned in multiple slices to initiate the Carving process in ilastik and connect the IAS across the image layers. Markers were further added if initial segmentation was deemed insufficient after proofreading. Manual correction was necessary for axonal branching, mitochondria attached to the myelin sheath, and the transition from myelinated to unmyelinated axonal segments.

The total processing time was about 2 weeks.

The comparison of segmentations from ilastik (Fig. 2a) and RaW (Fig. 2b) is based on the Jaccard index and the Sørensen-Dice index computed for individual IAS segmentations, and the foreground-restricted Rand F-score (V^rand^) and the information theoretic F-score (V^info^) (Arganda-Carreras et al. 2015) computed for IAS segmentations of all axons. The Jaccard index for each axon (Fig. 2d) is the pixel number of the intersection divided by the pixel number of the union of IAS segmentations from both methods. The Sørensen-Dice index for each axon (Fig. 2e) is the pixel number of the intersection divided by the average pixel number of IAS segmentations from both methods. The foreground-restricted rand F-score and the information theoretic F-score are closely related to the Rand index and the variation of information, respectively (Arganda-Carreras et al. 2015).

**Fig. 2.**
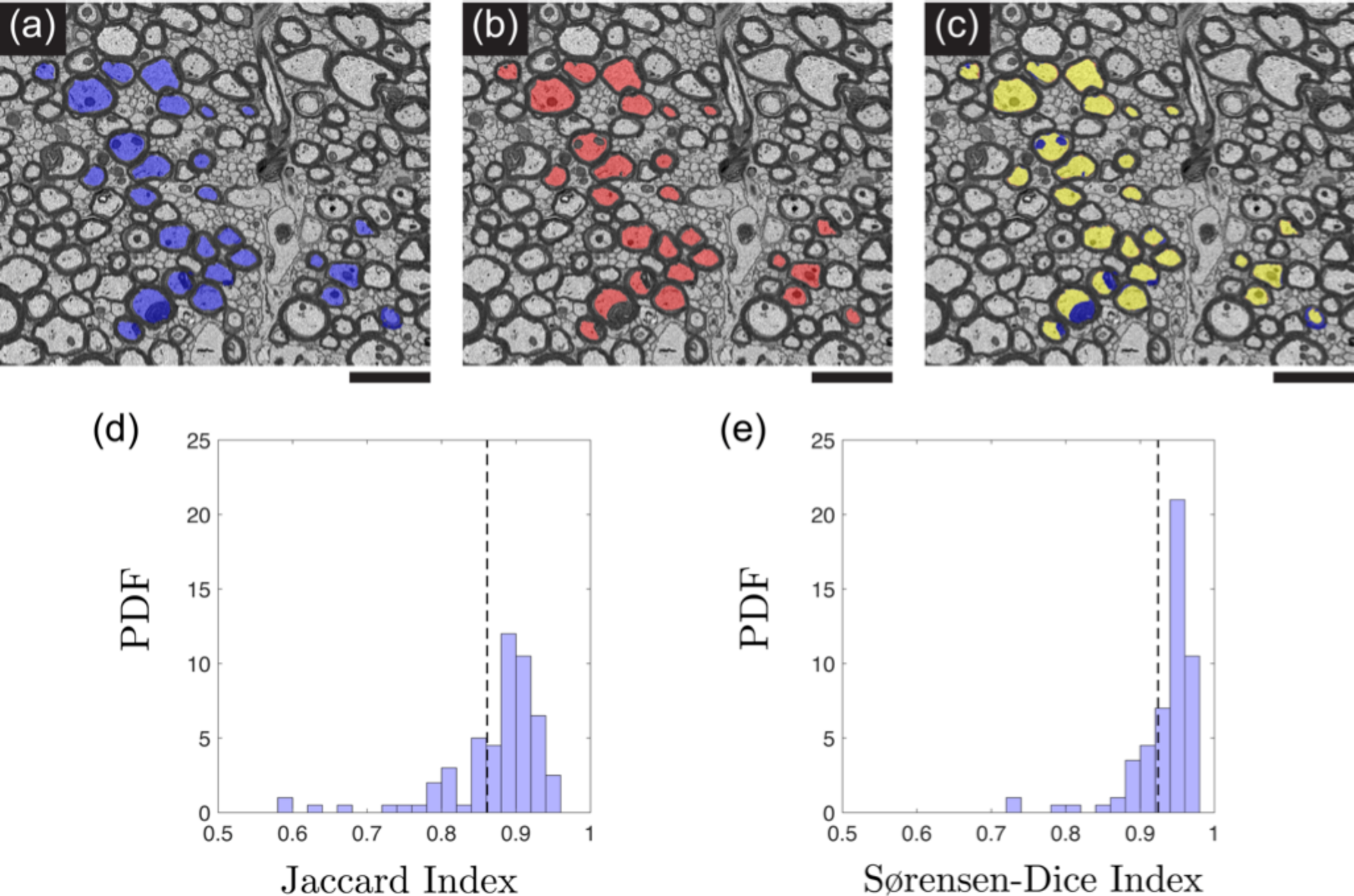
IAS segmented by (a) ilastik, Carving (blue pixels), (b) RaW (red pixels) and (c) both methods (intersection, yellow pixels) in a representing slice. The histogram of the (d) Jaccard index and the (e) Sørensen-Dice index for the comparison of IAS segmentations from the two methods. The scale bar below (a-c) is 4 μm

### Myelin sheath

Each axon’s individual myelin sheath (Fig. 1e) was obtained by overlapping the myelin mask and the expanded IAS segmentation (Kleinnijenhuis et al. 2017), which is dilated by a myelin thickness upper bound = 0.4 μm, a biologically plausible value for myelinated axons in the brain WM (see *Discussions, limitations* for further explanations). Cases of adjacent axons with touching myelin sheath are segmented by applying a non-weighted distance transform and watershed on the binary mask including all of the segmented IAS.

### Fiber Orientation Distribution (FOD)

For the segmented IAS of each axon, the corresponding axon skeleton connecting all centers of mass in each slice was computed. To mimic the microstructure coarse-grained by diffusion (Novikov et al. 2016; Novikov et al. 2018a), the piecewise linear skeleton of each segmented axon was then smoothed by a Gaussian kernel with a variance *σ^2^ = L^2^/4* (Novikov et al. 2014), where *L* is the diffusion length 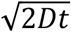, where *D* = 2 μm^2^/ms is a typical value for the intrinsic intra-axonal diffusion coefficient (Novikov et al. 2018b; Veraart et al. 2016), and *t* = [1, 10, 100] ms is the diffusion time range as applicable for dMRI.

### Dispersion angle

The dispersion angle of axon segments was calculated by using definitions, corresponding to *2d* histological observations and dMRI measurements.

In histology, each microscopy image is a *2d* cross-section of 3*d* structures. To compare our results with previous *2d* histological studies, we projected all the axon segments onto a projection plane parallel to the bundle’s mean direction, and calculated the projected dispersion angle *θ*_*2d*_*(φ)*, defined by the standard deviation of angles between the projected segments and the mean bundle direction, along which we rotated the projection plane with an azimuthal angle φ.

On the other hand, the SH-based dMRI models (Jespersen et al. 2017; Novikov et al. 2018b) are sensitive to an effective dispersion angle 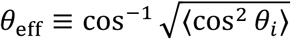, where the individual axon segment’s dispersion angle θ_*i*_ is the angle between the axon segment’s direction and the bundle’s mean direction. The *θ*_*eff*_*(φ)* is calculated by only including axon segments oriented between φ *± Δ*φ/*2*, where *Δ*φ = 12°.

Furthermore, we express the FOD 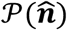 as a linear combination of SH basis 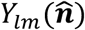 via

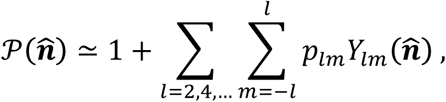

where *p*_*lm*_ is the SH coefficients. The rotational invariants *p*_*l*_ are determined by the 2-norm of SH coefficients via (Novikov et al. 2018b; Reisert et al. 2017)

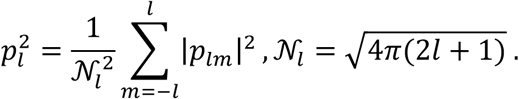

The normalization factor is chosen such that *p*_*0*_ *= l* and *p*_*1*_ *ϵ* [0, l] for *I >* 0. The FOD of axon segments is contributed only by even orders *l* (Novikov et al. 2018b; Reisert et al. 2017) since the FOD has an antipodal symmetry. The dispersion angle *θ*_*p2*_ estimated by rotational invariants is given by (Novikov et al. 2018b)

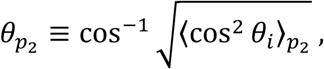

and

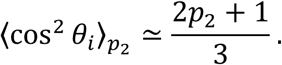

The value of *θ*_*p2*_ is theoretically close to the value of 0_eff_ since both of their definitions are related with the rms of cos θ_*i*_.

To simplify the relationship between rotational invariants and the order *l*, Reisert et al. proposed a simple Poisson kernel for the axial symmetric FOD with the form p_*l*_ = *λ*^*l*^, where *λ ϵ* [0,1] is a dispersion parameter (Reisert et al. 2017). We tested the applicability of this Poisson kernel, which effectively corresponds to the multipole expansion of a Coulomb potential.

### Inner axonal diameter

To evaluate the influence of the diffusion time on the axonal diameter distribution measured with dMRI, we aligned each axon’s main direction (denoted as z_axon_) parallel to the z-axis, cut off 1μm at both ends, in order to create the axon skeleton (a line connecting the center of mass of each slice), and calculated the cross-sectional area *Ω* for each slice perpendicular to the skeleton. Assuming axon as a circular cylinder, its inner diameter is defined as the diameter of an equivalent circle with the same area: 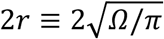.

### g-ratio

The g-ratio, manifesting axon myelination, was estimated as follows: we aligned the IAS and the myelin sheath parallel to the z-axis for each axon, cut off 1 μm at both ends, and estimated the outer diameter 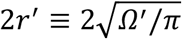, where the cross-sectional area *Ω’*, perpendicular to the axon skeleton, contains both the IAS and the myelin sheath. The g-ratio was estimated by 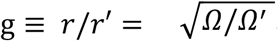.

## Results

### RaW and ilastik (Carving

The metrics to compare segmentations from ilastik and RAW for all segmented IAS are summarized in Table 1.

**Table 1.** Similarity metrics to compare IAS segmentations by ilastik and RaW

| Similarity metric | IAS segmentations (ilastik, RaW) |
| --- | --- |
| Jaccard Index | 0.86 |
| Sørensen-Dice index | 0.92 |
| $V^{\text{rand}}$ | 0.75 |
| $V^{\text{info}}$ | 0.90 |

In Fig. 2a, the IAS segmented using ilastik covers all contents in IAS, such as the cytoplasm and organelles (including mitochondria and nuclei attached to the myelin sheath by manual corrections). However, Fig. 2b shows that the IAS segmented by using RaW fails to cover organelles attached to the myelin sheath since these structures are always considered as part of the myelin mask when applying a bi-Gaussian model for the thresholding.

Fig. 2d-e shows the Jaccard index and the S0rensen-Dice index to compare the IAS segmentations from the two methods (ilastik and RaW). For most axons, both indices are high, manifesting the robustness of our random-hopping segmentation pipeline; this is also demonstrated by high values of similarity metrics shown in Table 1.

### FOD

Fig. 3a-b shows each axon’s smoothed skeleton displayed with a 3d view angle (Fig. 3a) or a 2d projection (Fig. 3b) for *t* = [1, 10, 100] ms. Longer diffusion time leads to a longer diffusion length and a wider smoothing kernel, and effectively smooths out each axon’s tortuous skeleton. The FOD, displayed either on a triangulated spherical surface (Womersley 2017) (Fig. 3c) or by a SH-constructed 3d glyph up to the order of *l* = 10 (Politis 2016) (Fig. 3d), indicates that longer diffusion time corresponds to a narrower fiber dispersion, which can be quantified by the dispersion angle shown in Fig. 4.

**Fig. 3.**
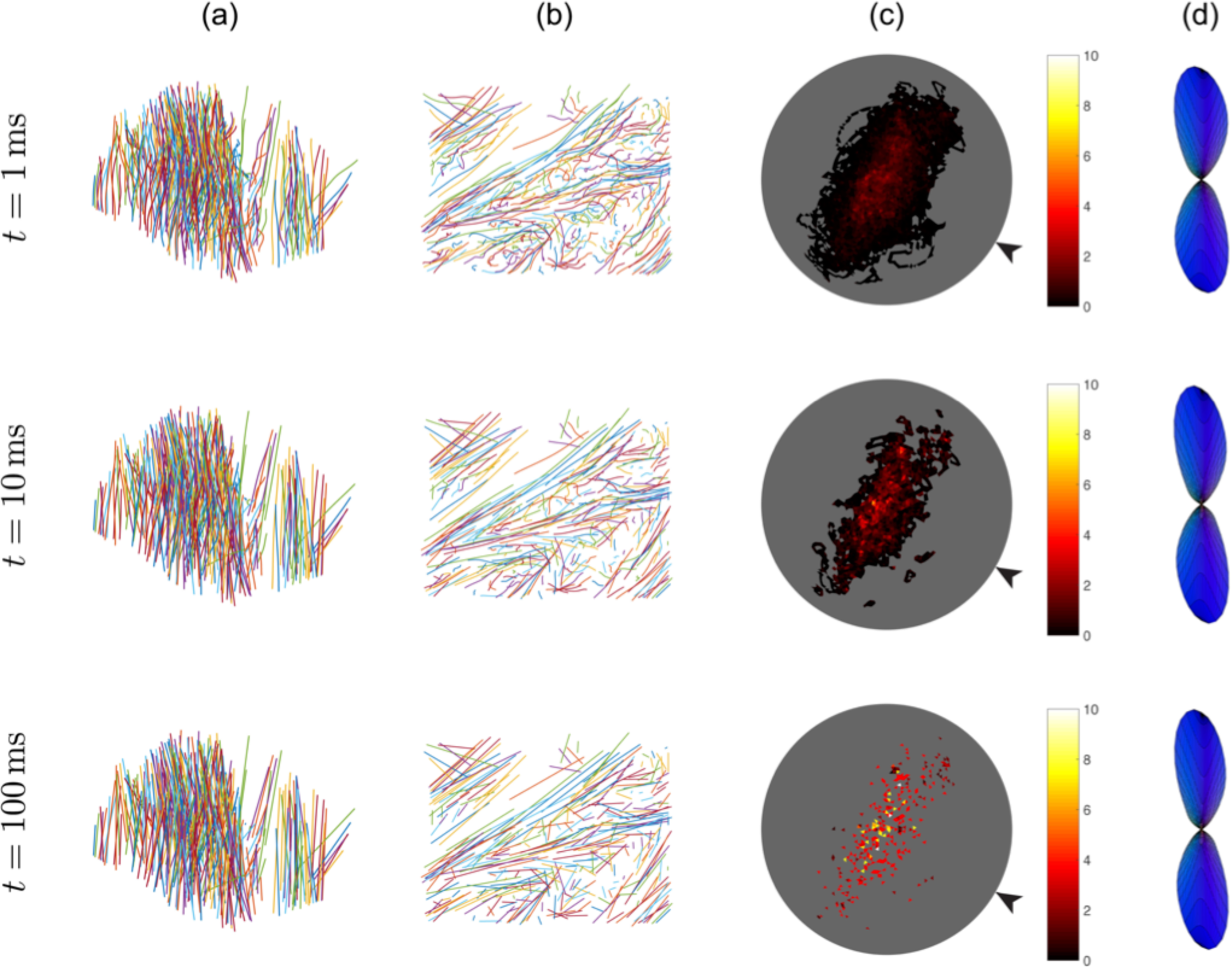
(a) The skeleton of each segmented axon is smoothed to mimic the diffusion time-dependent coarse-grained microstructure along each axon’s main direction with diffusion time *t* = [1, 10, 100] ms. (b) The skeleton of each segmented axon in (a) was viewed from another view angle. Each axon becomes effectively straighter for longer diffusion times. (c) The FOD of tangent vectors of all axon segments, starting at the center of a unit sphere, shows the intrinsic axonal dispersion. The unit of the colorbar is steradian^−1^. (d) The 3*d* FOD glyph was generated by fitting the FOD in (c) to spherical harmonics up to the order of *l* = 10. Arrows in (c) indicate the view angle for FOD glyphs in (d)

To compare with the previous study from (Schilling et al. 2016) that applies a smoothing kernel of a 1 μm width, we also fitted the FOD of *t* = 1 ms *(σ* = 1 μm) to a Bingham distribution (Bingham 1974; Sotiropoulos et al. 2012), yielding fitting parameters *k*_*1*_ = 22.2 and *k_2_* = 4.6. The orientation dispersion index, defined by (Mollink et al. 2017)

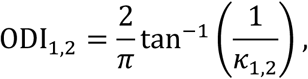

is ODI1 = 0.029 and ODI2 = 0.136.

### Dispersion angle and rotational invariants

Fig. 4a-b shows the (cross-sectional) projected dispersion angle θ_2*d*_(0) (Fig. 4a) and the effective dispersion angle θ_eff_(φ) (Fig. 4b) with respect to the azimuthal angle φ for *t* = [1, 10, 100] ms. Generally, θ_2*d*_(φ) varies between 8° to 23°, and θ_eff_(φ) varies between 5° to 31°.

**Fig. 4.**
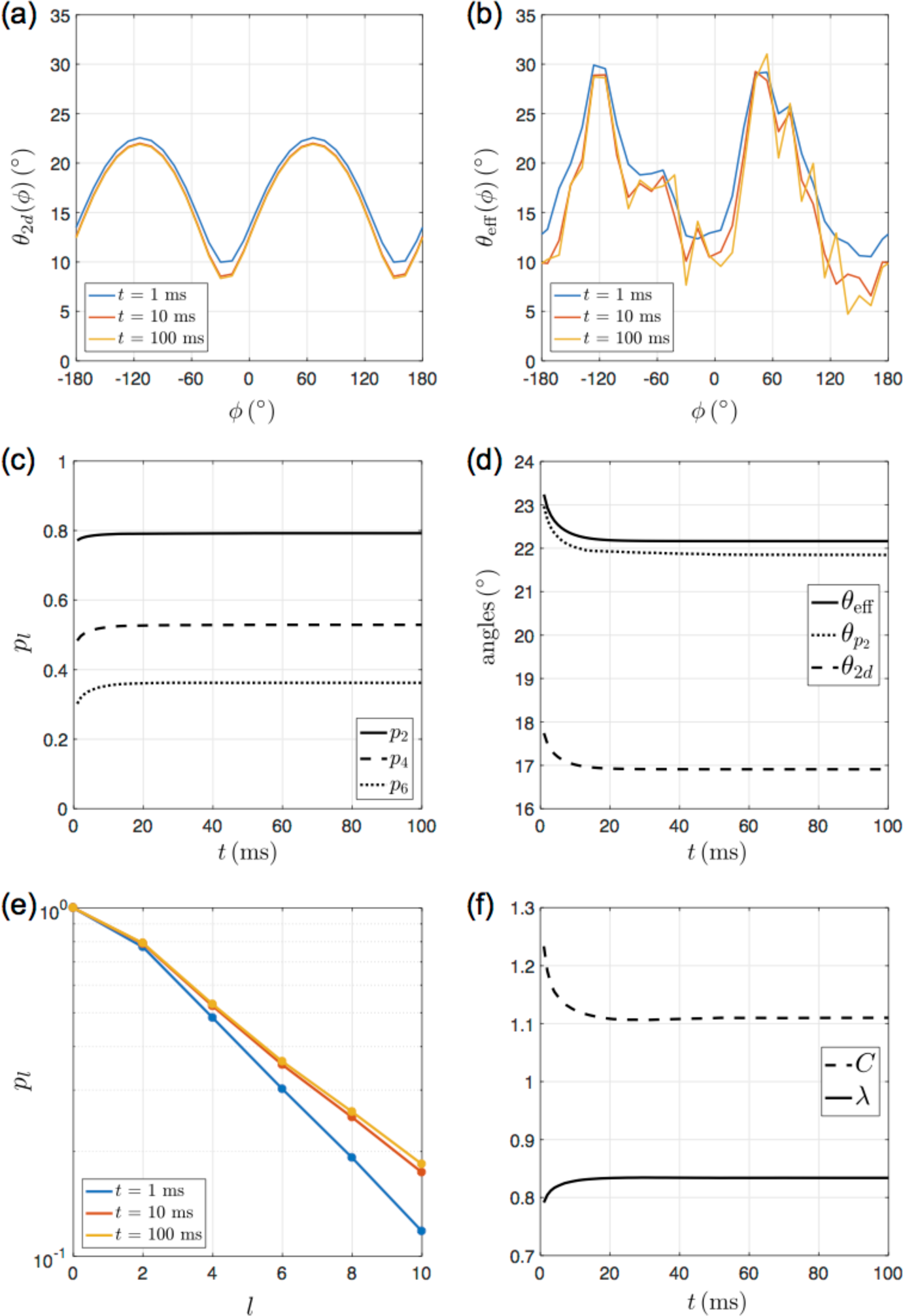
(a) Projected dispersion angle θ_2d_(φ) and (b) dMRI-sensitive dispersion angle θ _eff_(φ) (calculated within a bin width Δ φ = 12°) with respect to the azimuthal angle 0 at diffusion time *t* = [1, 10, 100] ms. (c) The rotational invariants (p_2_, p_4_, p_6_) show a time-dependence < 0.4 % for diffusion time *t* = 20-100 ms. (d) The dispersion angle averaged over all φ shows a time-dependence of ~ 1° for diffusion time *t* = 1-100 ms. (e) Rotational invariants *p*_*l*_ with respect to the even orders *l*= 2, 4, …, 10 at diffusion time *t* = [1, 10, 100] ms. (f) Dispersion parameters of the modified power-law relation *(λ,C* in Eq. (1)) obtained by using a linear fit of logp_A_ with respect to *l*= 2-10 for diffusion time *t* = 1-100 ms

In Fig. 4c, the time-dependence of rotational invariants is small for *p*_*2*_ (3 %), moderate for p_4_ (9 %), and large for *p*_*6*_ (20 %) for *t* ranging over 1-20 ms, and is insignificant (< 0.4%) for p_2_, p_4_ and p_6_ for diffusion time > 20 ms.

To further display the time-dependence of the dispersion angle, we calculated the averaged dispersion angle of the three definitions for *t* ranging over 1-100 ms in Fig. 4d, where the averaged *θ*_*2d*_ (rms of *θ* _*2d*_(φ) at φ = 1°, 2°, …,360°) decreases with the diffusion time, from 17.7° *(t* = 1 ms, *σ* = 1 μm) to 16.9° *(t* = 100 ms, *σ* = 10 μm); the dMRI-sensitive dispersion angle *θ*_*e*__ff_ (calculated by using all axon segments), demonstrates a similar time-dependence, and is always larger than the corresponding histology-observed projected dispersion angle *θ*_*2d*_ for all diffusion times; the dispersion angle *θ*_*p2*_, estimated by rotational invariants of the FOD, is slightly smaller than *θ*_*eff*_ and shows a similar time-dependence as well. Generally, the time-dependence of dispersion angles *(θ_2d_, θ_eff_, θ_p2_*) is small (~ 1° for *t* = 1-100 ms) and negligible for diffusion time 20 ms.

In Fig. 4e, rotational invariants are plotted as a function of even orders *l*, and seem to obey a power-law over the range *l* = 2, 4, …, 10. The base of the power-law is estimated via the slope log λ, and the predicted p_0_ is given by the intercept *C* at *l* = 0 (Eq. (1) in *Discussions).* By definition, p_0_ ≡ 1; yet, this power-law relation predicts a p_0_ > 1, as manifested by an intercept *C* 1 at *I* = 0. In Fig. 4f, the time-dependence of dispersion parameters is small for λ (5 %) and moderate for *C* (11%) for *t* ranging over 1-100 ms.

### g-ratio

The histogram of the outer axonal diameter, inner axonal diameter, and the g-ratio are shown in Fig. 5a-c. Their mean ± SD is for the outer axonal diameter = 1.68 ± 0.46 μm, inner axonal diameter = 1.03 ± 0.41 μm, and g-ratio = 0.59 ± 0.09. For further comparisons with other studies, we also reported the median and the interquartile range (IQR in parenthesis) for the outer axonal diameter = 1.61 (0.52) μm, inner axonal diameter = 0.95 (0.52) μm, and g-ratio = 0.60 (0.12).

**Fig. 5.**
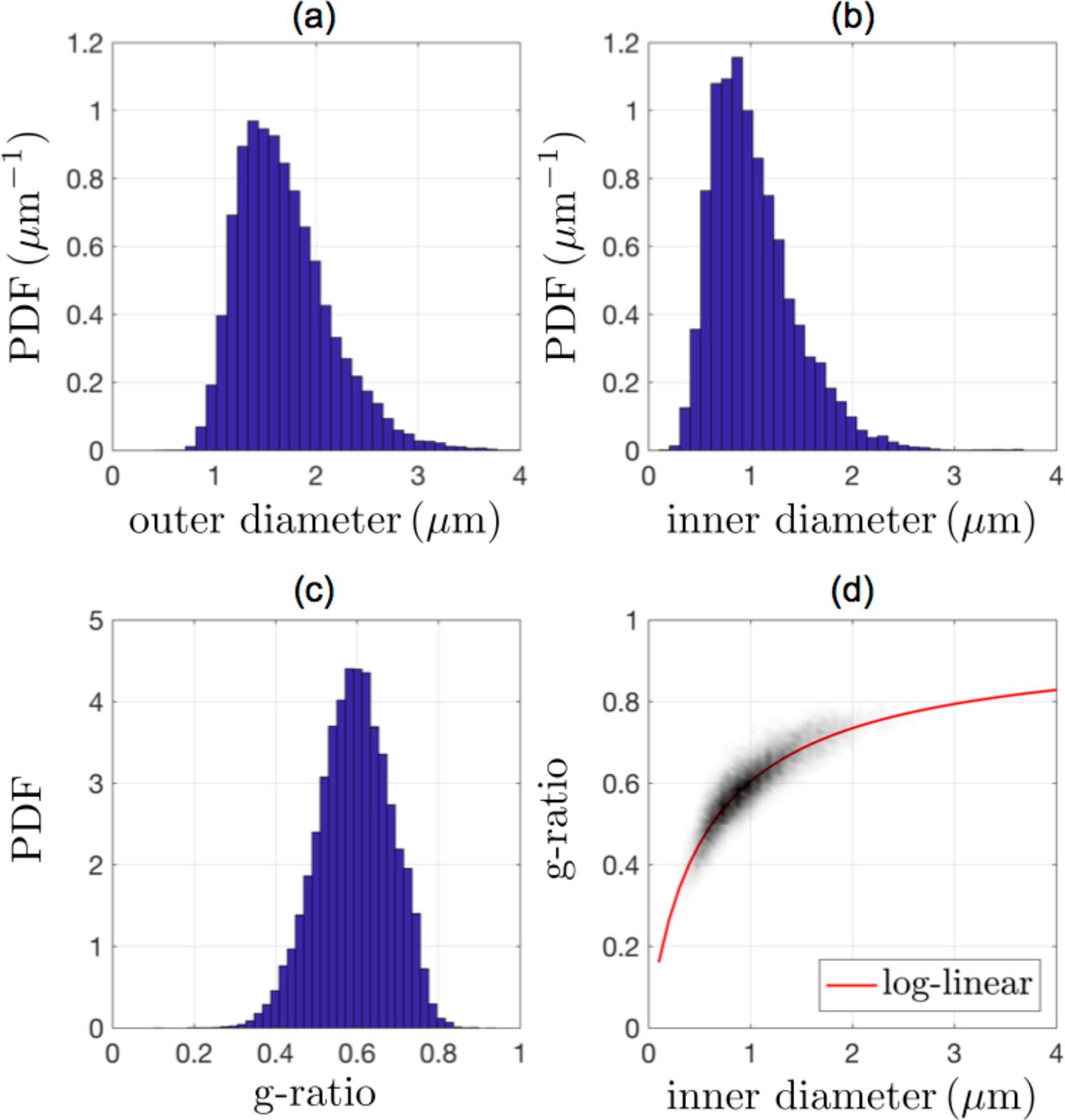
The histogram of (a) outer axonal diameter, (b) inner axonal diameter, and (c) genuine g-ratio. The relation of g-ratio and inner diameter is shown in (d) as a 2*d* histogram, fitted by the log-linear equation (red curve) proposed by (Berthold et al. 1983)

Fig. 5d shows the dependency of g-ratio on the inner diameter. This relationship is consistent with previous studies and fitted well by the reported log-linear (Berthold et al. 1983; Little and Heath 1994; West et al. 2015), where the myelin sheath thickness is proportional to the number of myelin lamellae *nl = C_0_+C_1_ ·*(2*r*) + *C*_*2*_*·* ln(2*r*). The red curve in Fig. 5d is the fit, and corresponding parameters are *C*_*0*_*k* = 0.31 μm, *C*_*1*_*k* = 0.02, and *C*_*2*_*k* = 0.02 μm, where *k* is the myelin lamellar width.

### Inner axonal diameter

Fig. 6a-c shows the inner axonal diameter variation along each axon, smoothed by a Gaussian kernel with the variance *σ*^*2*^ for *t* = [1, 10, 100] ms, along with the corresponding diameter histogram (Fig. 6d). Fig. 6e shows the time-dependences of the average diameter 2⟨*r*⟨ and dMRI-sensitive effective diameter *2r*_*eff*_, where 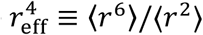 (Burcaw et al. 2015) based on the signal attenuation in the wide-pulse limit (Neuman 1974).

**Fig. 6.**
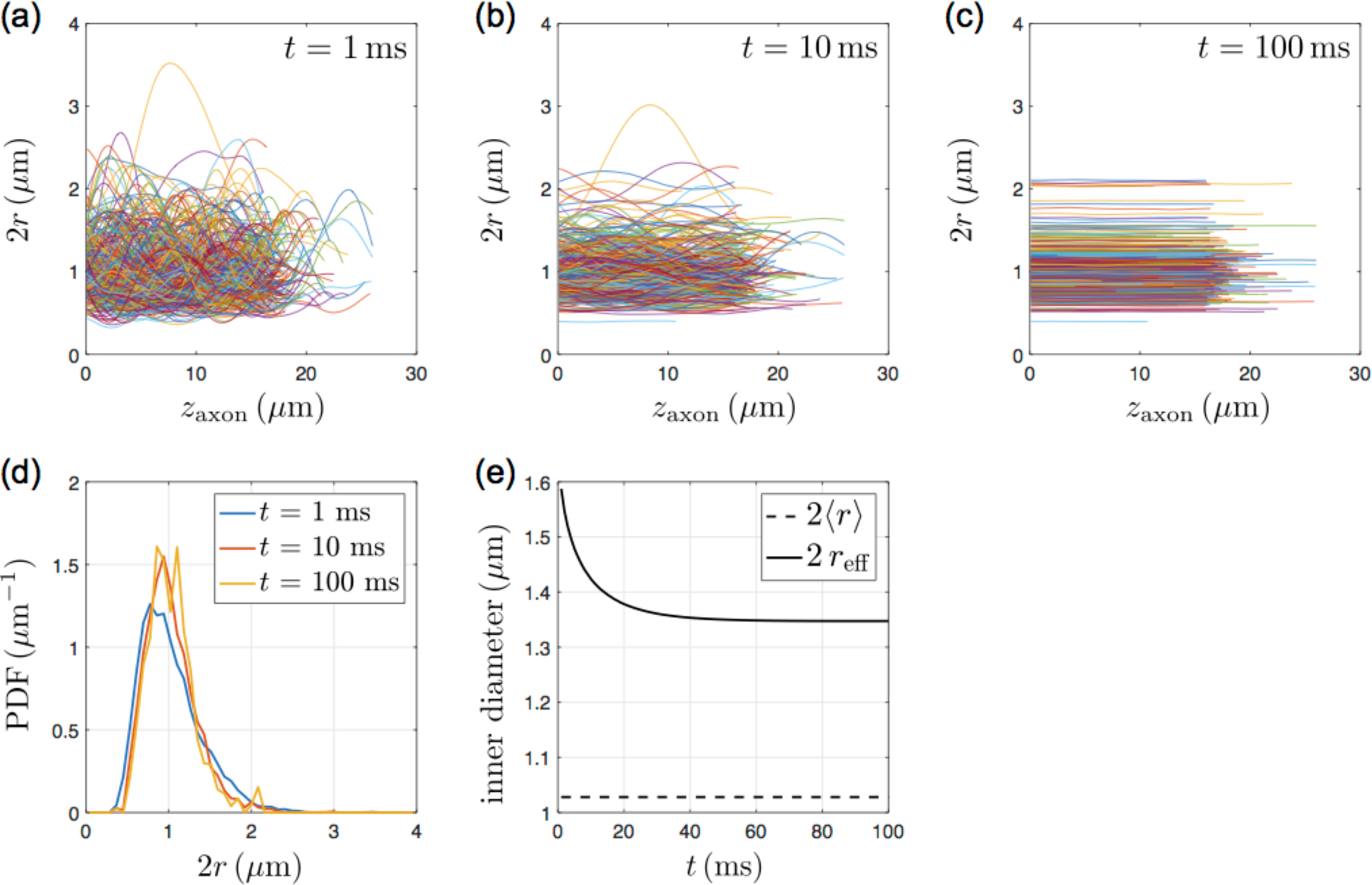
The axonal diameter (2*r*) variation, estimated by the cross-sectional area perpendicular to the skeleton and displayed along the main direction of each axon (z_axon_), was smoothed by a Gaussian kernel mimicking the diffusion process with an effective diffusion time *t* = (a) 1 ms, (b) 10 ms, and (c) 100 ms. (d) The diameter histogram becomes narrower with longer diffusion time. (e) The average diameter 2⟨*r*⟨ has no significant time-dependence, whereas the dMRI-sensitive effective diameter 2*r*_eff_, where 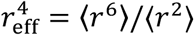 (Burcaw et al. 2015), has a non-trivial time-dependence for diffusion time *t* < 50 ms

At short diffusion time (Fig. 6a), the axonal diameter varies a lot within each axon, whereas, at long diffusion time (Fig. 6c), axonal diameter variation is smoothed out within each axon. Therefore, the axonal diameter distribution at short diffusion time (blue curve in Fig. 6d) is slightly wider than that at long diffusion time (yellow curve in Fig. 6d). In Fig. 6e, the mean diameter 2⟨*r*⟨ shows no obvious time-dependence. However, the dMRI-sensitive effective diameter 2*r*_eff_ shows significant time-dependence, ~ 18% change, for *t* ranging over 1-100 ms.

Table 2 shows the inner axonal diameter estimations using different definitions, such as the short and long axis length of the fitted ellipse with the same second-moments of the axon cross-section, and the inscribed circle diameter calculated by employing the distance transform to the axon cross-section. The median of the inscribed circle diameter provides the smallest diameter estimates = 0.71 μm, and, in contrast, the mean of the long axis length provides the largest diameter estimates = 1.24 μm. The cross-section perpendicular to the skeleton has an average eccentricity of 0.63 ± 0.15 (mean ± SD) and 0.65 (0.21) (median (IQR)), indicating that axons are in general approximately elliptical, rather than circular or cylindrical (Abdollahzadeh et al. 2017).

**Table 2.** Inner axonal diameter of myelinated axons, calculated by the equivalent circle diameter (cross-sectional area), the short axis length and long axis length of the fitted ellipse, and the inscribed circle diameter. Standard deviation (SD) and interquartile range (IQR) are shown in the parenthesis

| | Inner axonal diameter ($\mu\text{m}$) | |
| --- | --- | --- |
|  | Mean (SD) | Median (IQR) |
| Equivalent circle diameter = $2\sqrt{\Omega/\pi}$ | 1.03 (0.41) | 0.95 (0.52) |
| Short axis length | 0.92 (0.38) | 0.84 (0.47) |
| Long axis length | 1.24 (0.50) | 1.14 (0.64) |
| Inscribed circle diameter | 0.76 (0.33) | 0.71 (0.41) |

## Discussion

We successfully segmented ~ 51,000 axon segments from ~ 300 myelinated axons by using a semi-automatic random-hopping-based algorithm. Diffusion time-dependence of the orientation-related and size-related tissue characteristics of the brain white matter microstructure is, for the first time, analyzed via *3d* high-resolution EM images. The estimated dispersion angle of myelinated axons has negligible diffusion time-dependence at typical diffusion times observed with dMRI. In contrast, the estimated inner axonal diameter has a nontrivial time-dependence at diffusion times relevant for both pre-clinical and clinical diffusion imaging.

Here, we discuss our RaW algorithm as compared to a commonly used interactive segmentation tool, as well as how our results of orientation-related (FOD, dispersion angle, rotational invariants) and size-related (g-ratio, inner axonal diameter) tissue parameters compare to previous histological and MRI studies. Finally, we address some limitations of our methods.

### Segmentation methods (ilastik and RaW)

Although ilastik greatly facilitates tracing individual axons (one axon at a time), it is still labor-intensive. Each individual axon required many training data (object markers and background markers) in multiple slices, and the segmentation results change dynamically while new markers are added, increasing the loading of proofreading. The program was quite accurate in delineating the IAS, but more manual input was required at specific locations, such as axonal branching, mitochondria or nuclei attached to the myelin sheath.

In contrast, RaW algorithm is straightforward, and depends solely on the quality of the binary myelin mask. An imperfect myelin mask results in a segmentation (random-hopping trajectory) infiltrating into other axons or compartments, such as extra-axonal space. In this study, we successfully segmented ~ 70 % of axons crossing the central slice, albeit ~ 30 % of axons were deleted by the proofreading because of the leaky myelin mask. Random-hopping-based method minimizes the need of manual seeding and proofreading (1-2 days, ~ 300 axons/person), reducing hard labor and simplifying the segmentation pipeline, c.f. 2 weeks, ~ 100 axons/person by using ilastik (Carving). The manual seeding step can be further automated by extracting the regional maxima from the distance transform map of the myelin mask and the dilated edges (Abdollahzadeh et al. 2017). Confirmed with segmentation results by the Carving function of ilastik, RaW algorithm is robust and reliable to segment the IAS of myelinated axons. Similar to ilastik, mitochondria attached to the myelin sheath are deemed to be part of the myelin mask and therefore not delineated accurately, though it should be possible to separately identify the mitochondria using the semi-automatic pipeline incorporating superpixel-based simple-linear-iterative-clustering (SLIC) method (Abdollahzadeh et al. 2017; Achanta et al. 2012).

For machine-learning-based segmentation methods, it is time-consuming to produce training, development, and test data set from 3d EM data. Using RaW method, we can rapidly generate enough data to train and validate other segmentation algorithms. Also, simultaneously acquiring EM and light microscopy images (Wouters and Koerten 1982) provides extra features by staining specific organelles, facilitating automatic seeding for RAW and potentially model training data for machine-learning-based methods.

### FOD, dispersion angle, and rotational invariants

Our study presents a 3d EM-based extraction of fiber dispersion in the mouse brain genu of CC, and reports good agreement with dispersion estimated using confocal microscopy and light microscopy. In particular, the FOD at *t* = 1 ms (smoothed by *σ* = 1 μm) fitted to a Bingham distribution suggests *a k*_1_ value (≈ 22), corresponding to small dispersion (e.g., single fiber dispersion), consistent with existing literature ~ 21 (Schilling et al. 2016), where the structure tensor analysis was applied to 3d stacks of confocal microscopy images in monkey brains by using a Gaussian kernel (standard deviation = 1 μm) to calculate spatial derivatives. In contrast, the *k*_2_ value (≈ 5 in this study), related to large dispersion (e.g., fiber fanning/branching), is different from the previous study ~ 12 (Schilling et al. 2016), probably influenced by the sampling site in CC.

In addition, the estimated dispersion angle is in agreement with previous histological studies yielding a dispersion of ~ 18.1° for the human brain CC in (Ronen et al. 2014) and 17° for the rat brain CC in (Leergaard et al. 2010). Remarkably, it is also in agreement with the recently dMRI-estimated in vivo fiber dispersion ~ 17° in major human WM tracts (Veraart et al. 2016). Along directions with small fiber spread (φ ≈ −30° and 150° in Fig. 4a-b), the dispersion angle is ~ 8° with ODI_2_ = 0.029, consistent with values estimated from confocal microscopy images of the CC (Schilling et al. 2018).

Furthermore, the estimated dispersion angle is comparable between dMRI studies using different diffusion times, as we found the time-dependence of dispersion angles is minimal (~ 5 *%* for *t* = 1-100 ms in Fig. 4d). Similarly, for diffusion time > 20 ms, the time-dependence of rotational invariants (*p*_2_,*p*_4_, *p*_6_) is minimal (< 0.4 % for *t* = 20-100 ms in Fig. 4c), assuring the assumption of the time-independence of rotational invariants in SH-based models (Novikov et al. 2018b; Reisert et al. 2017).

As initially proposed by (Reisert et al. 2017), we verified that the rotational invariants approximately obey a power-law (Poisson kernel) of the order *l* for *l* = 2-10, and found that this power-law behavior is not well-normalized and overshoots at *l* = 0 (Fig. 4e). To compensate for that, a negative isotropic term needs to be introduced into the power-law, losing the simplicity of the Poisson kernel, i.e.

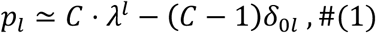

where *C* is a constant ≥ 1, and δ_0*l*_ is a Kronecker delta. Indeed, in Fig. 4f, the dispersion parameters (λ, C) are obtained by using a linear fit of log_p*l*_ with respect to *l* =2-10, whereby the fitted *C* (≈ 1.1-1.2) > 1 indicates the overshoot of the power-law relation (λ ≈ 0.8) at *l* = 0.

### g-ratio

We obtained a relatively smaller histology g-ratio value ≈ 0.6, as compared to previous histological studies: 0.808 for > 18-week-old mice (Mason et al. 2001), 0.81 for adult mice (West et al. 2015) (age not specified), and 0.76 for 2-month-old mice (Yang et al. 2016). This could be caused by (1) the difference of changes in myelin structures during the EM processing (Kirschner and Hollingshead 1980), such as fixation and dehydration, and (2) potentially inaccurate segmentation of the myelin sheath. Since we only segmented some axons, instead of all, the watershed algorithm cannot avoid overestimation of the segmented myelin sheath touching the myelin sheath of the unsegmented axons, leading to an overall overestimated myelin sheath thickness and a slightly underestimated g-ratio in our study.

Different definitions of g-ratio are used in the field of histology versus MRI, with the latter (gMRi) sometimes being called the aggregate g-ratio, as defined by 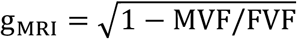 (Stikov et al. 2011), where MVF and FVF are the myelin volume fraction and the fiber volume fraction, respectively. However, while MRI models typically assume a single g-ratio value for the axons within an MRI voxel, we reported that the genuine g-ratio g from histology has a non-negligible variation over our sample size (Fig. 5d) which is much smaller than a typical MRI voxel (by an order of 100). In order to compare MRI measurement with histology, (West et al. 2016) proposed the following relation between the g_MRI_ and the genuine histology g:

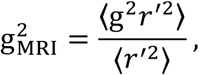

which corresponds to an estimated g_MRI_ = 0.64 in this study, in agreement with the aggregate g-ratio = 0.62 for rat brain CC in (Abdollahzadeh et al. 2017) and aggregate g-ratio = 0.69 for macaque brain CC in (Stikov et al. 2015).

### Inner axonal diameter and its distribution

By calculating the cross-sectional area perpendicular to the axon skeleton (Abdollahzadeh et al. 2017), the estimated inner axonal diameter is 1-2 *%* smaller than the estimation calculated by using the cross-sectional area perpendicular to axon’s main direction (data not shown). In other words, diameter estimations are not significantly affected by either considering axon skeletons or not as in our sample, since axons in CC are generally straight (mean sinuosity = 1.04).

Although the Gamma distribution is the most commonly used model for the inner axonal diameter distribution, the generalized extreme value distribution describes the diameter distribution better (Sepehrband et al. 2016a). This argument is also true in our data for the inner diameter distribution shown in Fig. 5b (fits to distributions are shown in *Appendix A* and Fig. 8).

In this study, we used the equivalent circle diameter to evaluate inner and outer axonal diameters and the genuine g-ratio. Alternatively, the inner axonal diameter can be estimated by other definitions, such as short and long axis length of the fitted ellipse, and the inscribed circle diameter (Table 2). Generally, median diameters are smaller than mean diameters by 6-9 %; compared with the equivalent circle diameter, the short axis length is smaller by ~11 %, the long axis length is larger by ~ 20 %, and the inscribed circle diameter is smaller by ~ 24 %. The inscribed circle diameter was used as a diameter estimate in many histology studies (Aboitiz et al. 1992; Caminiti et al. 2009; Liewald et al. 2014), while some studies, on the other hand, did not mention their calculation methods of inner diameters.

For the mouse brain CC, our inner diameter estimates (equivalent circle diameter) is larger than values reported in previous EM studies by a factor of 1.2-1.8: 0.47 μm for 45-day-old mice in (Sturrock 1980), 0.88 μm for > 18-week-old mice in (Mason et al. 2001), 0.56 μm for adult mice in (West et al. 2015) (age not specified), and 0.54 μm for an 8-week-old mouse in (Sepehrband et al. 2016a). This is potentially caused by differences in calculation methods (e.g., equivalent circle diameter, short axis length, inscribed circle diameter), the image quality, and the tissue shrinkage during fixation (Bozzola and Russell 1999).

The MRI-sensitive effective diameter *2r*_*eff*_ varies from 1.59 μm to 1.35 μm for *t* = 1-100 ms, values that are slightly larger than reported values of *2r*_*eff*_ in a previous histological study (1.32 μm) (Sepehrband et al. 2016b). However, in previous MRI literature, the dMRI-measured diameter is larger than our *2r*_*eff*_ estimation by a factor of ≥ 1.4, even when applying very strong diffusion-sensitive gradients **|*g|*** ≤ 1350 mT/m (Sepehrband et al. 2016b). This discrepancy could be due to neglecting the diffusion time-dependence of the extra-axonal signal (Burcaw et al. 2015; De Santis et al. 2016; Lee et al. 2017), or potentially due to misinterpreting the extra-axonal signal change as the intra-axonal one. In particular, when applying strong diffusion gradients, the dMRI-measured diameter is further biased by neglecting the higher order ***|g|**^4^* corrections (Lee et al. 2017) to the intra-axonal model in (Neuman 1974).

### Limitations

The random-hopping-based segmentation method depends heavily on the quality of the myelin mask. Mitochondria and nuclei directly attached to the inner myelin border are recognized as part of the myelin mask and need to be separately identified by other algorithms (Abdollahzadeh et al. 2017; Achanta et al. 2012).

While segmenting the individual myelin sheath for each axon, we assigned an upper bound for the myelin thickness (Kleinnijenhuis et al. 2017). The upper bound has to be chosen carefully when proofreading since the g-ratio has a strong dependence on this tuning parameter (Fig. 7): A small upper bound leads to an under-segmented myelin sheath and a large g-ratio, and a large upper bound leads to an over-expanded myelin sheath and a small g-ratio. Determining the upper bound of the myelin thickness is crucial when evaluating the g-ratio and the actual myelin thickness, whereas this upper bound is usually not reported in other studies.

**Fig. 7.**
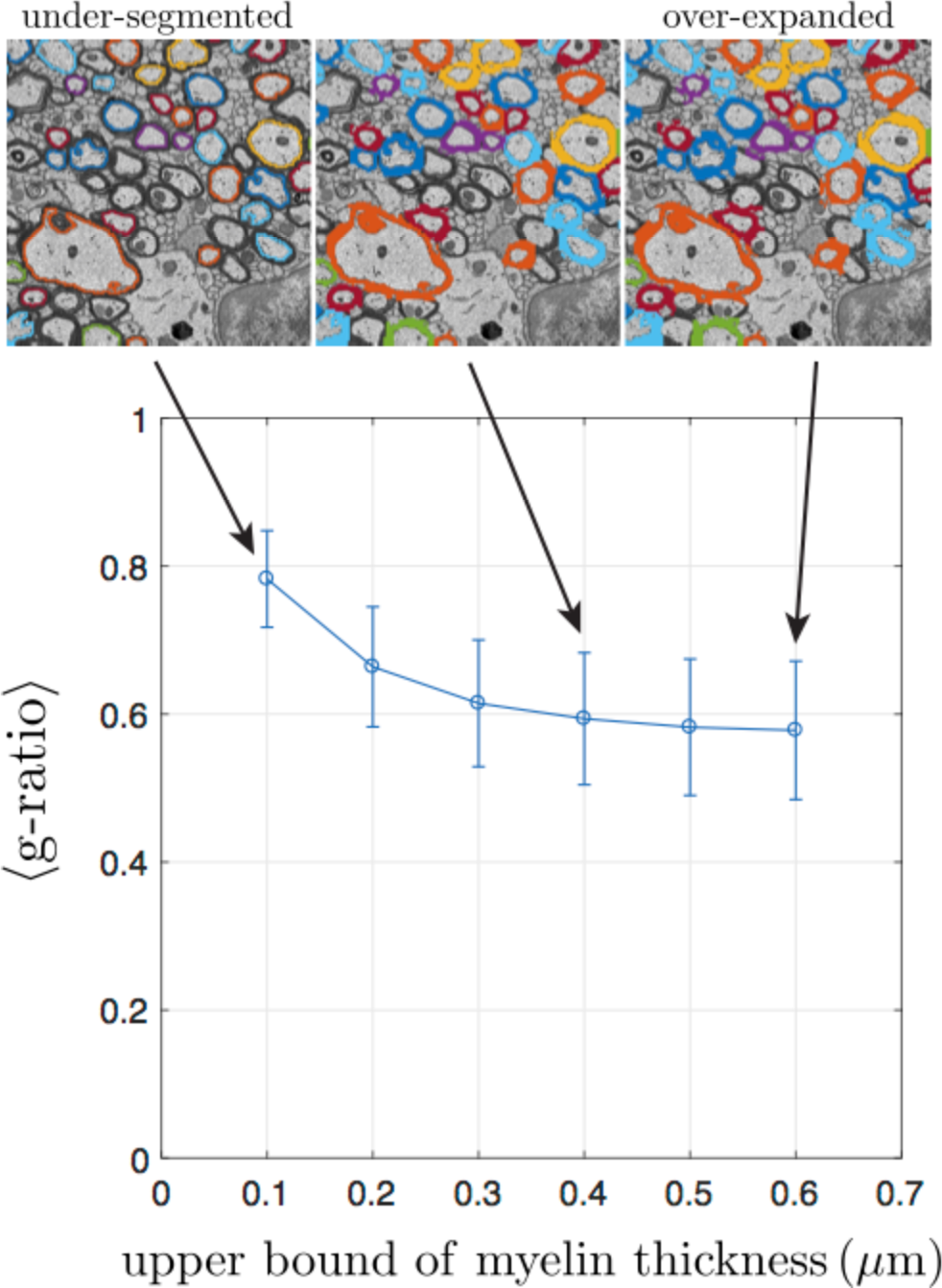
The artificial upper bound applied for myelin sheath segmentation influences the mean g-ratio. A small upper bound for the myelin thickness leads to under-segmented individual myelin sheaths (top left, upper bound = 0.1 μm). In contrast, a large upper bound causes overexpanded individual myelin sheaths (top right, upper bound = 0.6 μm). In this study, an upper bound of 0.3-0.4 μm results in appropriate individual myelin sheaths (top middle, upper bound = 0.4 μm)

Compared with other imaging techniques, such as light microscopy (Grussu et al. 2016; Ronen et al. 2014), polarized light imaging (Mollink et al. 2017), and confocal microscopy (Schilling et al. 2016; Schilling et al. 2018), the size of EM samples is relatively small, and results based on EM segmentations could be less representative because of the limited field-of-view.

In this study, we only focus on myelinated axons. Other structures, such as unmyelinated axons, astrocytes, and blood vessels, are also important and need to be segmented for a more comprehensive analysis.

## Conclusion

We present here a random-hopping-based segmentation method facilitating a 3*d* EM segmentation pipeline in brain white matter, with minimal labor for proofreading and manual seeding that could also be further automated. The 3*d* EM segmentation provides an accurate and reliable evaluation of the fiber orientation dispersion, and the calculated projected dispersion angle is compatible with previous 2*d* histological studies, as well as agreement of the estimated MRI-measured dispersion angle with previous MRI studies, with a very small diffusion time-dependence. Besides the fiber orientation information, the 3*d* EM segmentation enables to estimate the inner and outer axonal diameter as well as the g-ratio according to various definitions by analyzing the cross-section perpendicular to the axon skeleton. Our simulation shows that the diffusion time-dependence of the dMRI-derived axon diameter metric is nontrivial and has to be taken into account for studies mapping axonal diameters with dMRI.

## Acknowledgement

We would like to thank the NYULH DART Microscopy Lab Alice Liang, Kristen Dancel-Manning and Chris Patzold for their expertise in electron microscopy work, Kirk Czymmek and Pal Pedersen from Carl Zeiss for their assistance of 3*d* EM data acquisition, and Marios Georgiadis for the discussion of the myelin structure change caused by tissue preparations. Research was supported by the National Institute of Neurological Disorders and Stroke of the NIH under award number R21 NS081230 (Fieremans, E., Novikov, D. S., and Kim, S. G.) and R01 NS088040 (Fieremans, E. and Novikov, D. S.), and was performed at the Center of Advanced Imaging Innovation and Research (CAI2R, www.cai2r.net), an NIBIB Biomedical Technology Resource Center (NIH P41 EB017183, Fieremans, E., Novikov, D. S., and Kim, S. G.).

## Appendix A. Axonal diameter estimates in various definitions

In this section, distributions of axonal diameters are showed based on different definitions, such as equivalent circle diameter (Fig. 8a), short and long axis length of the fitted ellipse (Fig. 8b-c), and inscribed circle diameter (Fig. 8d). To compare with a previous study (Sepehrband et al. 2016a), we fitted the axonal diameter histogram to Gamma distribution and generalized extreme value (GEV) distribution in Fig. 8, which shows that GEV distribution fits better to the experimental diameter distribution (of all four definitions) than Gamma distribution does, consistent with the conclusion in (Sepehrband et al. 2016a). Also, GEV distribution has a longer tail than Gamma distribution does for thick axons in diameters > 3-5 μm, manifested by semi-logarithmic plots of diameter distributions in the bottom row of Fig. 8.

**Fig. 8.**
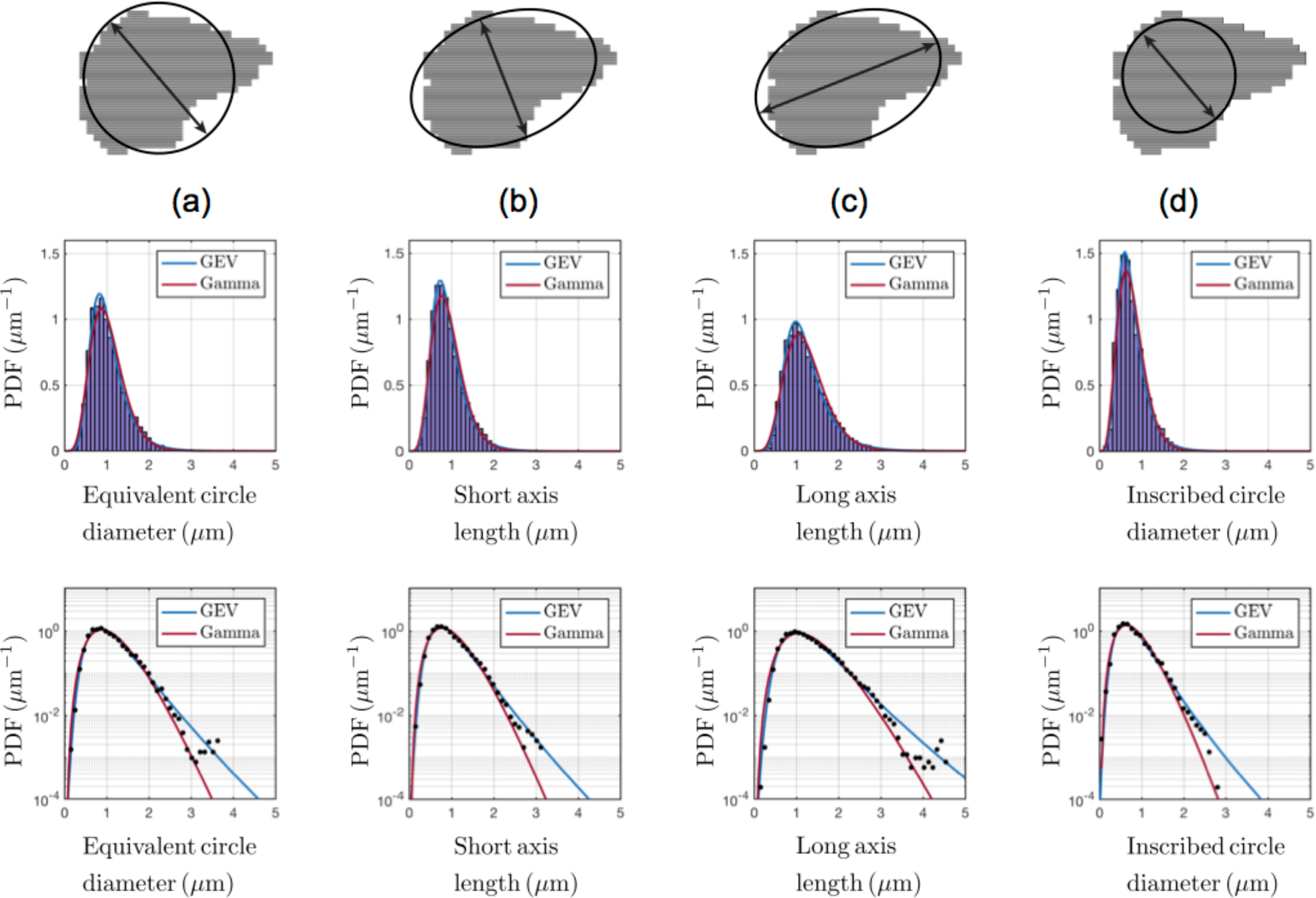
The distribution of axonal diameters, defined by (a) equivalent circle diameter calculated from the cross-sectional area, (b) short axis length and (c) long axis length of the fitted ellipse, and (d) Inscribed circle diameter. The upper row shows an exemplified axon cross-section (gray area) and the corresponding diameter estimates (double-arrowed lines). The middle row shows experimental diameter distributions (purple bars) and the fits based on the Gamma distribution (red) and the generalized extreme value distribution (GEV) (blue). The bottom row is the middle row displayed in a semi-logarithmic scale for experimental data (data points) and the fits (solid lines)

## Compliance with Ethical Standards

- Disclosure of potential conflicts of interest

1. Funding: This study was supported by the National Institute of Neurological Disorders and Stroke of the NIH under award number R21 NS081230 (Fieremans, E., Novikov, D. S., and Kim, S. G.) and R01 NS088040 (Fieremans, E. and Novikov, D. S.), and was performed at the Center of Advanced Imaging Innovation and Research (CAI2R, www.cai2r.net), an NIBIB Biomedical Technology Resource Center (NIH P41 EB017183, Fieremans, E., Novikov, D. S., and Kim, S. G.).
2. Conflict of Interest: The authors declare that they have no conflict of interest.
- Ethical approval: All procedures performed in studies involving animals were in accordance with the ethical standards of the institution or practice at which the studies were conducted. This article does not contain any studies with human participants performed by any of the authors.
- Informed consent: Not applicable.

## References

Abdollahzadeh A, Belevich I, Jokitalo E, Tohka J, Sierra A (2017) 3D Axonal Morphometry of White Matter bioRxiv:239–228

Aboitiz F, Scheibel AB, Fisher RS, Zaidel E (1992) Fiber composition of the human corpus callosum Brain Res 598:143–153

Achanta R, Shaji A, Smith K, Lucchi A, Fua P, Susstrunk S (2012) SLIC superpixels compared to state-of-the-art superpixel methods IEEE Trans Pattern Anal Mach Intell 34:2274–2282 doi:10.1109/TPAMI.2012.120

Adams R, Bischof L (1994) Seeded Region Growing Ieee T Pattern Anal 16:641–647

Alexander DC, Hubbard PL, Hall MG, Moore EA, Ptito M, Parker GJ, Dyrby TB (2010) Orientationally invariant indices of axon diameter and density from diffusion MRI Neuroimage 52:1374–1389 doi:10.1016/j.neuroimage.2010.05.043

Arganda-Carreras I et al. (2015) Crowdsourcing the creation of image segmentation algorithms for connectomics Front Neuroanat 9:142 doi:10.3389/fnana.2015.00142

Assaf Y, Basser PJ (2005) Composite hindered and restricted model of diffusion (CHARMED) MR imaging of the human brain Neuroimage 27:48–58 doi:10.1016/j.neuroimage.2005.03.042

Assaf Y, Blumenfeld-Katzir T, Yovel Y, Basser PJ (2008) AxCaliber: a method for measuring axon diameter distribution from diffusion MRI Magn Reson Med 59:1347–1354 doi:10.1002/mrm.21577

Barazany D, Basser PJ, Assaf Y (2009) In vivo measurement of axon diameter distribution in the corpus callosum of rat brain Brain 132:1210–1220 doi:10.1093/brain/awp042

Baron CA, Kate M, Gioia L, Butcher K, Emery D, Budde M, Beaulieu C (2015) Reduction of Diffusion-Weighted Imaging Contrast of Acute Ischemic Stroke at Short Diffusion Times Stroke 46:2136–2141 doi:10.1161/STR0KEAHA.115.008815

Benjamini D, Komlosh ME, Holtzclaw LA, Nevo U, Basser PJ (2016) White matter microstructure from nonparametric axon diameter distribution mapping Neuroimage 135:333–344 doi:10.1016/j.neuroimage.2016.04.052

Berthold CH, Nilsson I, Rydmark M (1983) Axon diameter and myelin sheath thickness in nerve fibres of the ventral spinal root of the seventh lumbar nerve of the adult and developing cat J Anat 136:483–508

Bingham C (1974) Antipodally Symmetric Distribution on Sphere Ann Stat 2:1201–1225

Bozzola JJ, Russell LD (1999) Electron microscopy: principles and techniques for biologists. Jones & Bartlett Learning,

Budde MD, Frank JA (2010) Neurite beading is sufficient to decrease the apparent diffusion coefficient after ischemic stroke Proc Natl Acad Sci U S A 107:14472–14477 doi:10.1073/pnas.1004841107

Burcaw LM, Fieremans E, Novikov DS (2015) Mesoscopic structure of neuronal tracts from time-dependent diffusion Neuroimage 114:18–37 doi:10.1016/j.neuroimage.2015.03.061

Caminiti R, Ghaziri H, Galuske R, Hof PR, Innocenti GM (2009) Evolution amplified processing with temporally dispersed slow neuronal connectivity in primates Proc Natl Acad Sci U S A 106:19551–19556 doi:10.1073/pnas.0907655106

De Santis S, Jones DK, Roebroeck A (2016) Including diffusion time dependence in the extra-axonal space improves in vivo estimates of axonal diameter and density in human white matter Neuroimage 130:91–103 doi:10.1016/j.neuroimage.2016.01.047

Dorkenwald S, Schubert PJ, Killinger MF, Urban G, Mikula S, Svara F, Kornfeld J (2017) Automated synaptic connectivity inference for volume electron microscopy Nat Methods 14:435–442 doi:10.1038/nmeth.4206

Duval T et al. (2015) In vivo mapping of human spinal cord microstructure at 300mT/m Neuroimage 118:494–507 doi:10.1016/j.neuroimage.2015.06.038

Grussu F, Schneider T, Yates RL, Zhang H, Wheeler-Kingshott C, DeLuca GC, Alexander DC (2016) A framework for optimal whole-sample histological quantification of neurite orientation dispersion in the human spinal cord J Neurosci Methods 273:20–32 doi:10.1016/j.jneumeth.2016.08.002

Jeong WK, Beyer J, Hadwiger M, Vazquez A, Pfister H, Whitaker RT (2009) Scalable and interactive segmentation and visualization of neural processes in EM datasets IEEE Trans Vis Comput Graph 15:1505–1514 doi:10.1109/TVCG.2009.178

Jespersen SN, Olesen JL, Hansen B, Shemesh N (2017) Diffusion time dependence of microstructural parameters in fixed spinal cord Neuroimage doi:10.1016/j.neuroimage.2017.08.039

Kaynig V et al. (2015) Large-scale automatic reconstruction of neuronal processes from electron microscopy images Med Image Anal 22:77–88 doi:10.1016/j.media.2015.02.001

Kirschner DA, Hollingshead CJ (1980) Processing for electron microscopy alters membrane structure and packing in myelin J Ultrastruct Res 73:211–232

Kleinnijenhuis M, Johnson E, Mollink J, Jbabdi S, Miller K (2017) A 3D electron microscopy segmentation pipeline for hyper-realistic diffusion simulations, ISMRM 25th annual meeting, Hawaii, USA Proceedings of the ISMRM annual meeting 25:1090

Komlosh ME, Ozarslan E, Lizak MJ, Horkayne-Szakaly I, Freidlin RZ, Horkay F, Basser PJ (2013) Mapping average axon diameters in porcine spinal cord white matter and rat corpus callosum using d-PFG MRI Neuroimage 78:210–216 doi:10.1016/j.neuroimage.2013.03.074

Lee H-H, Fieremans E, Novikov DS (2017) What dominates the time dependence of diffusion transverse to axons: Intra- or extra-axonal water? NeuroImage doi:https://doi.org/10.1016/j.neuroimage.2017.12.038

Leergaard TB, White NS, de Crespigny A, Bolstad I, D’Arceuil H, Bjaalie JG, Dale AM (2010) Quantitative histological validation of diffusion MRI fiber orientation distributions in the rat brain PLoS One 5:e8595 doi:10.1371/journal.pone.0008595

Li P, Murphy TH (2008) Two-photon imaging during prolonged middle cerebral artery occlusion in mice reveals recovery of dendritic structure after reperfusion J Neurosci 28:11970–11979 doi:10.1523/JNEUROSCI.3724-08.2008

Liewald D, Miller R, Logothetis N, Wagner HJ, Schuz A (2014) Distribution of axon diameters in cortical white matter: an electron-microscopic study on three human brains and a macaque Biol Cybern 108:541–557 doi:10.1007/s00422-014-0626-2

Little GJ, Heath JW (1994) Morphometric analysis of axons myelinated during adult life in the mouse superior cervical ganglion J Anat 184 (Pt 2):387–398

Mason JL, Langaman C, Morell P, Suzuki K, Matsushima GK (2001) Episodic demyelination and subsequent remyelination within the murine central nervous system: changes in axonal calibre Neuropathol Appl Neurobiol 27:50–58

Mollink J et al. (2017) Evaluating fibre orientation dispersion in white matter: Comparison of diffusion MRI, histology and polarized light imaging Neuroimage 157:561–574 doi:10.1016/j.neuroimage.2017.06.001

Neuman C (1974) Spin echo of spins diffusing in a bounded medium The Journal of Chemical Physics 60:4508–4511

Novikov DS, Jensen JH, Helpern JA, Fieremans E (2014) Revealing mesoscopic structural universality with diffusion Proc Natl Acad Sci U S A 111:5088–5093 doi:10.1073/pnas.1316944111

Novikov DS, Jespersen SN, Kiselev VG, Fieremans E (2016) Quantifying brain microstructure with diffusion MRI: Theory and parameter estimation arXiv preprint arXiv:161202059

Novikov DS, Kiselev VG, Jespersen SN (2018a) On modeling Magn Reson Med 79:3172–3193 doi:10.1002/mrm.27101

Novikov DS, Veraart J, Jelescu IO, Fieremans E (2018b) Rotationally-invariant mapping of scalar and orientational metrics of neuronal microstructure with diffusion MRI Neuroimage doi:10.1016/j.neuroimage.2018.03.006

Politis A (2016) Microphone array processing for parametric spatial audio techniques Reisert M, Kellner E, Dhital B, Hennig J, Kiselev VG (2017) Disentangling micro from mesostructure by diffusion MRI: A Bayesian approach Neuroimage 147:964–975 doi:10.1016/j.neuroimage.2016.09.058

Ronen I, Budde M, Ercan E, Annese J, Techawiboonwong A, Webb A (2014) Microstructural organization of axons in the human corpus callosum quantified by diffusion-weighted magnetic resonance spectroscopy of N-acetylaspartate and post-mortem histology Brain Struct Funct 219:1773–1785 doi:10.1007/s00429-013-0600-0

Salo RA, Belevich I, Manninen E, Jokitalo E, Grohn O, Sierra A (2018) Quantification of anisotropy and orientation in 3D electron microscopy and diffusion tensor imaging in injured rat brain Neuroimage 172:404–414 doi:10.1016/j.neuroimage.2018.01.087

Schilling K, Janve V, Gao Y, Stepniewska I, Landman BA, Anderson AW (2016) Comparison of 3D orientation distribution functions measured with confocal microscopy and diffusion MRI Neuroimage 129:185–197 doi:10.1016/j.neuroimage.2016.01.022

Schilling KG, Janve V, Gao Y, Stepniewska I, Landman BA, Anderson AW (2018) Histological validation of diffusion MRI fiber orientation distributions and dispersion Neuroimage 165:200–221 doi:10.1016/j.neuroimage.2017.10.046

Sepehrband F, Alexander DC, Clark KA, Kurniawan ND, Yang Z, Reutens DC (2016a) Parametric Probability Distribution Functions for Axon Diameters of Corpus Callosum Front Neuroanat 10:59 doi:10.3389/fnana.2016.00059

Sepehrband F, Alexander DC, Kurniawan ND, Reutens DC, Yang Z (2016b) Towards higher sensitivity and stability of axon diameter estimation with diffusion-weighted MRI NMR Biomed 29:293–308 doi:10.1002/nbm.3462

Shepherd GM, Raastad M, Andersen P (2002) General and variable features of varicosity spacing along unmyelinated axons in the hippocampus and cerebellum Proc Natl Acad Sci U S A 99:6340–6345 doi:10.1073/pnas.052151299

Sommer C, Straehle C, Koethe U, Hamprecht FA Ilastik: Interactive learning and segmentation toolkit. In: Biomedical Imaging: From Nano to Macro, 2011 IEEE International Symposium on, 2011. IEEE, pp 230–233

Sotiropoulos SN, Behrens TE, Jbabdi S (2012) Ball and rackets: Inferring fiber fanning from diffusion-weighted MRI Neuroimage 60:1412–1425 doi:10.1016/j.neuroimage.2012.01.056

Stikov N et al. (2015) In vivo histology of the myelin g-ratio with magnetic resonance imaging Neuroimage 118:397–405 doi:10.1016/j.neuroimage.2015.05.023

Stikov N, Perry LM, Mezer A, Rykhlevskaia E, Wandell BA, Pauly JM, Dougherty RF (2011) Bound pool fractions complement diffusion measures to describe white matter micro and macrostructure Neuroimage 54:1112–1121 doi:10.1016/j.neuroimage.2010.08.068

Sturrock RR (1980) Myelination of the mouse corpus callosum Neuropathol Appl Neurobiol 6:415–420

Sun D, Roth S, Black MJ Secrets of optical flow estimation and their principles. In: Computer Vision and Pattern Recognition (CVPR), 2010 IEEE Conference on, 2010. IEEE, pp 2432–2439

Sun D, Roth S, Black MJ (2014) A quantitative analysis of current practices in optical flow estimation and the principles behind them International Journal of Computer Vision 106:115–137

Tang-Schomer MD, Johnson VE, Baas PW, Stewart W, Smith DH (2012) Partial interruption of axonal transport due to microtubule breakage accounts for the formation of periodic varicosities after traumatic axonal injury Exp Neurol 233:364–372 doi:10.1016/j.expneurol.2011.10.030

Tariq M, Schneider T, Alexander DC, Gandini Wheeler-Kingshott CA, Zhang H (2016) Bingham-NODDI: Mapping anisotropic orientation dispersion of neurites using diffusion MRI Neuroimage 133:207–223 doi:10.1016/j.neuroimage.2016.01.046

Veraart J, Fieremans E, Novikov DS (2016) Universal power-law scaling of water diffusion in human brain defines what we see with MRI arXiv preprint arXiv:160909145

West KL, Kelm ND, Carson RP, Does MD (2015) Quantitative analysis of mouse corpus callosum from electron microscopy images Data Brief 5:124–128 doi:10.1016/j.dib.2015.08.022

West KL, Kelm ND, Carson RP, Does MD (2016) A revised model for estimating g-ratio from MRI Neuroimage 125:1155–1158 doi:10.1016/j.neuroimage.2015.08.017

Wilke SA et al. (2013) Deconstructing complexity: serial block-face electron microscopic analysis of the hippocampal mossy fiber synapse J Neurosci 33:507–522 doi:10.1523/JNEUR0SCI.1600-12.2013

Womersley RS (2017) Efficient spherical designs with good geometric properties arXiv preprint arXiv:170901624

Wouters CH, Koerten HK (1982) Combined light microscope and scanning electron microscope, a new instrument for cell biology Cell Biol Int Rep 6:955–959

Yang HJ, Vainshtein A, Maik-Rachline G, Peles E (2016) G protein-coupled receptor 37 is a negative regulator of oligodendrocyte differentiation and myelination Nat Commun 7:10884 doi:10.1038/ncomms10884

Zaimi A, Wabartha M, Herman V, Antonsanti PL, Perone CS, Cohen-Adad J (2018) AxonDeepSeg: automatic axon and myelin segmentation from microscopy data using convolutional neural networks Sci Rep 8:3816 doi:10.1038/s41598-018-22181-4

Zhang H, Schneider T, Wheeler-Kingshott CA, Alexander DC (2012) NODDI: practical in vivo neurite orientation dispersion and density imaging of the human brain Neuroimage 61:1000–1016 doi:10.1016/j.neuroimage.2012.03.072

